# A new species of *Travassosinema* (Oxyuridomorpha: Travassosinematidae) from an Indian millipede, *Trigoniulus corallinus* (Gervais)

**DOI:** 10.1101/2020.08.02.232728

**Authors:** Somnath Bhakat

**Affiliations:** Dept. of Zoology, Rampurhat College, Rampurhat – 731224, Dist. Birbhum, West Bengal, India

**Keywords:** Bengal, gamopetallous, lateral alae, nematode, standard length, tail length

## Abstract

*Travassosinema bengalensis* n. sp. is described from the hind gut of the spirobolid millipede, *Trigoniulus corallinus* (Gervais) from West Bengal, India. Females of the new species differ from the only known Indian species, *T. travassosi* Rao, 1958 by several characters namely tail length, length of oesophagous, size of egg, extension of umbraculum etc. It is very similar to other three species of *Travassosinema, T. travassosi, T. thyropygi* Hunt, 1996 and *T. claudiae* Morffe & Hasegawa, 2017 as all of them lack lateral alae and body contraction posterior to vulva. Except *T. claudiae*, it differs from all other species from millipedes by longest tail length (60% SL) and differs from *T. claudiae* by shorter oesophagous length and location of vulva.

A new method for presentation of morphometric data (in percentage to standard length) in nematode is suggested.

On the basis of phylogenetic analysis, it is suggested that umbraculum bearing genera, *Indiana, Pulchrocephala* should be excluded from the family Travassosinematidae.

## Introduction

Among nematode parasites of millipede, *Travassosinema* Rao, 1958 is the most peculiar genus with a characteristic structure of six cephalic umbraculum which is absent in other known genera like *Thelastoma, Rhigonema, Deukemia* etc. Rao (1958) first described the genus and a species of *Travassosinema, T. travassoei* from a millipede, *Spirostreptus* sp. from Hyderabad, India. Later, more number of species was described from African, Asian and Australisian millipedes viz. *T. dechambrieri* Adamson, 1987; *T. morobecola* Hunt, 1993; *T. sulawesiense* Hunt, 1993; *T. thyropygi* Hunt, 1996 and *T. claudiae* Morfee and Hasegawa, 2017. Except millipede three other species of *Travassosinema* are recorded from other invertebrate host namely *T. mirabile* Spiridonov and Ivanova, 1998 from earthworm, T. jaidenae Jex, Schneider, Rose and Cribb, 2005 from wood-burrowing cockroach and *T. dalei* Spiridonov and Cribb, 2012 from a larvae of scarabaeid beetle. This paper deals with the second species of the genus *Travassosinema*, next to *T. travassoei* Rao, 1958 from Indian millipede.

## Materials and methods

Several specimens of *Trigoniulus corallinus* (Gervais) (Diplopoda: Spirobolida) were collected by hand from road side at Suri, Birbhum district, West Bengal, India for nematode study. In such attempt, three specimens of the female nematode were found to contain *Travassosinema*. The first specimen of the female nematode was identified from the millipede collected on 23.VI. 20 (treated as holotype). Later other six specimens were collected from other two female millipedes along with the genera, *Thelastoma* and *Dudekemia* on 26.VI. 20. and 03. VII. 20.

Millipedes were dissected by severing the head and epiproct with a razor blade as described by Phillips et al. (2016). The whole gut was pulled out intact from the body cavity with a fine tip forcep and put in a watch glass containing saline water. The gut was sectioned in three parts: fore-, mid- and hind-gut (Crawford et al.1983). Each portion of gut was taken in a slide with a drop of saline water and tore with the help of a fine needle tip. Nematodes were removed from intestinal tissue carefully and sorted on the basis of general features. Specimens were killed and fixed in hot (60 – 70°C) 4% formalin, then processed to anhydrous glycerine via slow evaporation method (Seinhorst, 1959). The specimens were mounted on slides in a glycerine drop embedded in a nail polish ring.

Nematodes were measured with ocular micrometer in a compound microscope. Sketches were made with the help of camera lucida.

## Results

### Systematics

Family: Travassosinematidae Rao, 1958

Genus: *Travassosinema* Rao, 1958

*Travassosinema bengalensis* n. sp. (Fig. 1).

**Fig. 1.**
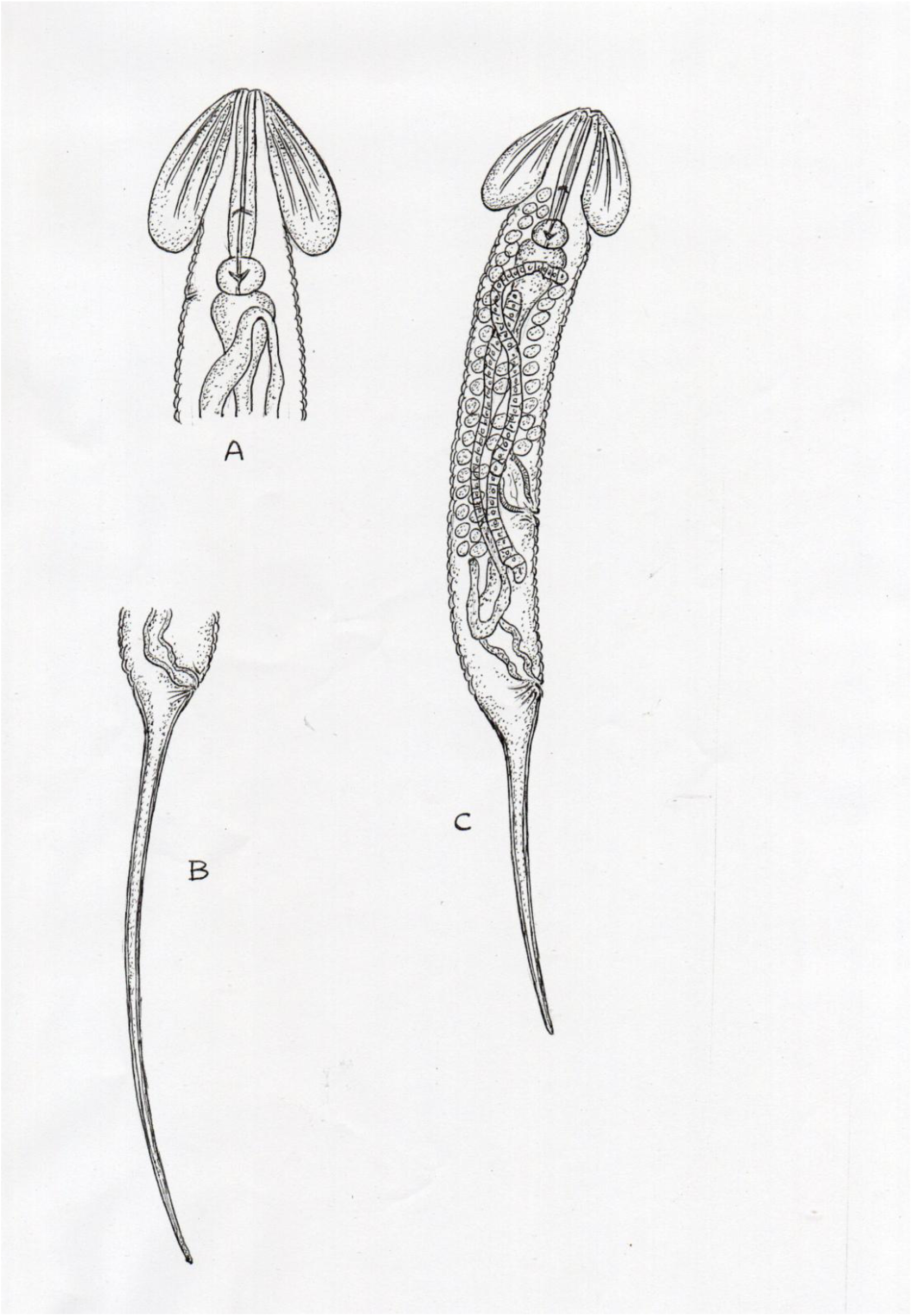
*Travassosinema bengalensis* n. sp. Female. A. Anterior region (1mm = 8.5 µm). B. Tail (1mm = 8.0µm) C. Habitus (1mm = 13.0µm (Lateral view).

### Type material

Holotype: Female (2.61 mm), Suri (87°32’00’’E, 23°55’ 00’’N), Birbhum district, West Bengal, India; in *Trigoniulus corallinus* (Diplopoda: Spirobolida); 23. VI. 2020; S. Bhakat collector; Zoological Museum, Dept. of Zoology, Rampurhat College. Rampurhat – 731224, Dist. Birbhum, West Bengal, India.

Paratype: Six females, other information are same as holotype.

### Description

Female. Small nematode is with cephalic umbraculum which looks like petals of a flower. Body more or less cylindrical with narrow umbraculum bearing region and a long filiform tail ending in a fine tip (60% of standard length). Body cuticle bears transverse annulations of about 7.5 µm width. Annulations begin at point where umbraculum joins with the body and extend to the level of anus. Lateral ala is totally absent.

Oral opening triradiate and is surrounded by three well developed partially overlapped lips, one dorsal and two subventrals. Umbraculum consists of six radially arranged and posteriorly directed alae. Each ala is a membranous expansion of body cuticle, anterior attached with the body and posterior free with rounded lobe and extending to the level of anterior to oesophageal bulb. The whole structure looks like gamopetallous corolla of China rose. Ala is widest in the middle with convex edge. Like other species each ala or ‘petal’ is reinforced by three rib like structure of different length.

Oesophagous is with narrow cylindrical procorpous. Oesophageal corpous not demarcated from isthmus and bulb. Bulb is slightly oval in shape with well developed valve plate. Anterior portion of intestine attached to bulb is broader. Nerve ring at the posterior 3/4^th^ region of the procorpous. Excretory pore ventral and near the base of oesophagous. Vulva is a small transverse slit posterior to midventral length of the body (60% level from anterior to standard length).

Genital tract is amphidelphic in nature. Body is not contracted posterior to vulva. Ovaries are slender. Anterior ovary reflexed and surrounded the intestine and may extend beyond the basal bulb anteriorly. Posterior ovary is reflexed beyond the junction of intestine with the rectum. Anterior uterus is with small rounded seminal receptacle. Eggs are oval in shape and may be arranged in three rows. In the holotype female, there are numerous eggs arranged in three rows in the middle of the body but in a single row in the anterior portion of the body, extending beyond the midregion of the procorpous. Morphometric data of *Travassosinema bengalensis* is presented in Table 1. (conventional form) and Table 2. (modified form).

**Table 1.**
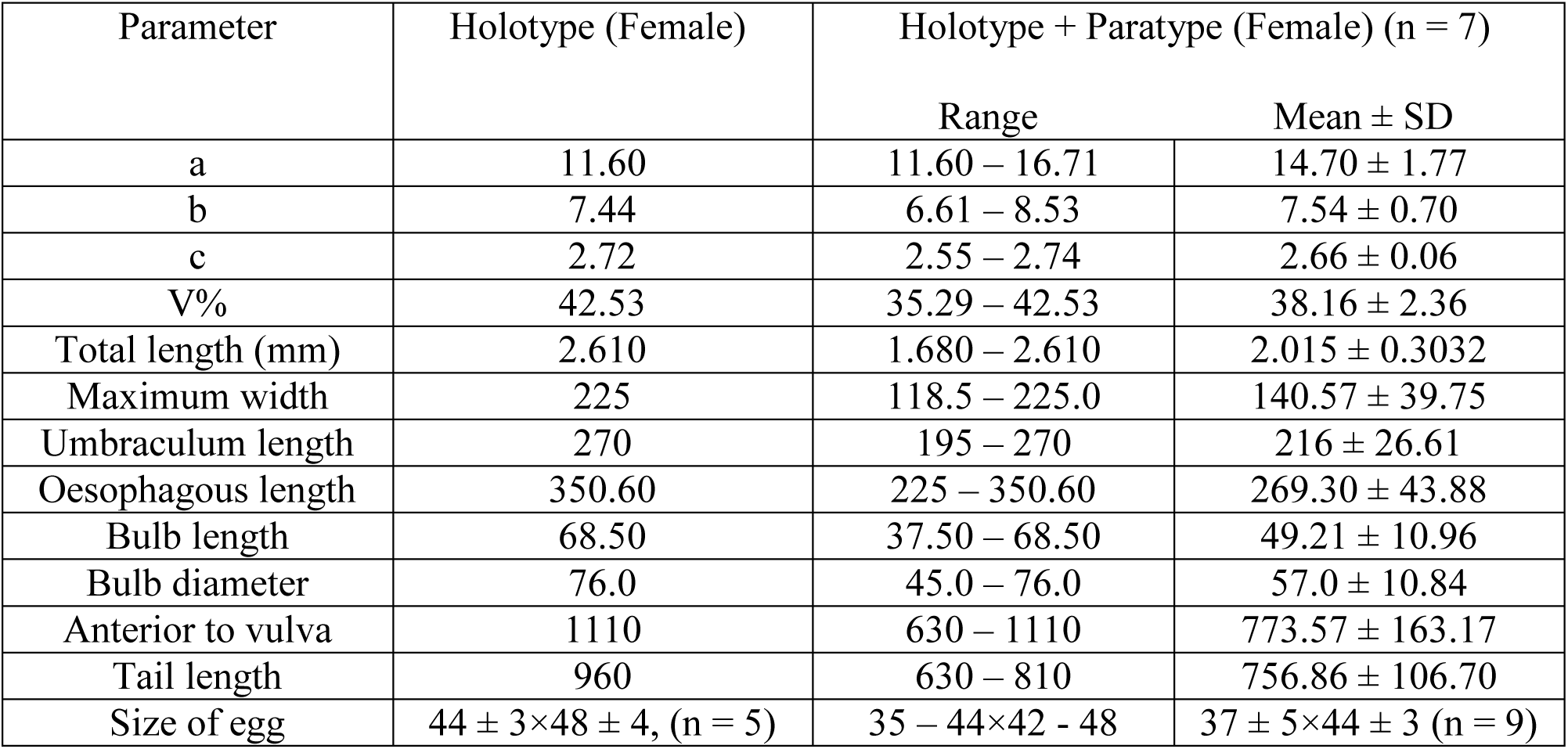
Morphometrics of *Travassosinema bengalensis* n. sp. (measurements are in µm unless otherwise indicated)

| Parameter | Holotype (Female) | Holotype + Paratype (Female) (n = 7) |  |
| --- | --- | --- | --- |
| | | Range | Mean ± SD |
| a | 11.60 | 11.60 – 16.71 | 14.70 ± 1.77 |
| b | 7.44 | 6.61 – 8.53 | 7.54 ± 0.70 |
| c | 2.72 | 2.55 – 2.74 | 2.66 ± 0.06 |
| V% | 42.53 | 35.29 – 42.53 | 38.16 ± 2.36 |
| Total length (mm) | 2.610 | 1.680 – 2.610 | 2.015 ± 0.3032 |
| Maximum width | 225 | 118.5 – 225.0 | 140.57 ± 39.75 |
| Umbraculum length | 270 | 195 – 270 | 216 ± 26.61 |
| Oesophagous length | 350.60 | 225 – 350.60 | 269.30 ± 43.88 |
| Bulb length | 68.50 | 37.50 – 68.50 | 49.21 ± 10.96 |
| Bulb diameter | 76.0 | 45.0 – 76.0 | 57.0 ± 10.84 |
| Anterior to vulva | 1110 | 630 – 1110 | 773.57 ± 163.17 |
| Tail length | 960 | 630 – 810 | 756.86 ± 106.70 |
| Size of egg | 44 ± 3 × 48 ± 4, (n = 5) | 35 – 44 × 42 – 48 | 37 ± 5 × 44 ± 3 (n = 9) |

**Table 2.** Morphometrics of *Travassosinema bengalensis* n. sp. (n=7) (modified form).

| Parameter | Holotype | Range (Mean ± SD) |
| --- | --- | --- |
| Total length (mm) | 2.610 | 1.680 – 2.610 (2.015 ± 0.30) |
| Standard length (SL) (mm) | 1.650 | 1.050 – 1.650 (1.258 ± 0.19) |
| %SL |  |  |
| Body width (BW) | 13.63 | 9.43 – 13.63 (11.05 ± 0.62) |
| Oesophagous length | 21.25 | 18.75 – 24.32 (21.47 ± 1.70) |
| Anterior to vulva | 67.27 | 56.76 – 67.27 (61.15 ± 3.44) |
| Bulb length | 4.15 | 3.45 – 4.54 (3.89 ± 0.26) |
| Bulb diameter | 4.61 | 4.27 – 5.08 (4.42 ± 0.37) |
| Umbraculum length | 16.36 | 14.77 – 20.27 (17.03 ± 1.46) |
| Tail length | 58.18 | 58.18 – 64.36 (60.31 ± 2.21) |
| Total length | 158.18 | 157.47 – 164.36 (160.31 ± 2.21) |
| Egg size (%BW) | 19.56 ± 2 × 21.33 ± 3 (n=5) | 19.56 – 21.90 × 21.33 – 28.57<br>(20.37 ± 2 × 25.58 ± 3) (n=9) |

Type locality: Suri (87°32’00’’E, 23°55’00’’N), Birbhum district, West Bengal, PIN 731224, India.

Type host: Female *Trigoniulus corallinus* (Gervais) (Diplopoda: Spirobolida). Site: Hindgut.

Etymology: Latin name genitive case meaning “bengal” in reference to its occurrence.

### Differential diagnosis

Different species of the genus *Travassosinema* can be distinguished by the presence or absence of lateral alae, body contraction posterior to vulva and relative size of the pharynx. The present species, *T. bengalensis*, resembles with only three species of *Travassosinema* from millipedes, viz. *T. travassosi, T. thyropygi* and *T. claudiae* as all of them lack lateral alae and contraction of posterior vulva. *T. bengalensis* differs from *T. travassosi* and *T. thyropygi* by longer tail (60% of SL vs. 46 – 55% in *T. travassosi* and 46% in *T. thyropygi*) though tail length of *T. claudiae* and *T. bengalensis* is almost same (60% of SL). In *T. bengalensis* eggs are smaller in size compared to other three species (44× 48 vs. > 44× 48). In the present species cephalic umbraculum extends anterior to oesophageal bulb but in *T. claudiae*, it is comparatively shorter, extends until posterior third of the procorpous and in *T. travassosi* it is longer, extends to midpoint of basal bulb. Compared to *T. claudiae* and *T. travassosi*, length of oesophagous in *T. bengalensis* is shorter (b = 7.54 vs. 8.21 in *T. claudiae* and 18 in *T. travassosi*). Furthermore, the vulva of *T. bengalensis* is more posteriorly located than in *T. claudiae* (V% = 35.92 – 42.53 vs. 32.53 – 38.18). Moreover, compare to all the reported six species of *Travassosinema* from millipede, *T. bengalensis* bears longest tail (60% vs. < 60% of SL).

### Key to seven species of *Travassosinema* from millipedes

1. Presence of lateral alae ……………………………… 2 -Absence of lateral alae …………………………… 3
2. Body length more than 3 mm …………………………… *T. sulawesiense* -Body length less than 3 mm …………………………… *T. morbecola*
3. Presence of contraction of the body posterior to the level of vulva …………………………… *T. dechambrieri* -Absence of contraction of the body posterior to the level of vulva …………………………… 4
4. Tail length more than 60% of standard length …………………………… *T. bengalensis* Tail length less than 60% of standard length …………………………… 5
5. Longer oesophagous length (b > 10) …………………………… *T. travassosi* -Shorter oesophagous length (b < 10) …………………………… 6
6. Nerve ring located in the procorpous-isthmus junction …………………………… *T. claudiae* -Nerve ring located anterior to basal bulb …………………………… *T. thyropygi*

## Discussion

The conventional method to present morphometric data of nematode in actual value is not rational, because actual data never depicts proportion or percentage of any body parts in relation to its body length. So it is more scientific to present size of different body parts in proportion to its body length. In nematode, tail length of different genera or species is also variable and may be often genera or species specific. As for example, in *Dudekemia*, the tail is very short (5 – 6%), in *Rhigonema*, it is 7 – 9%, in *Thelastoma*, tail is medium in size (16 – 19%) but in *Stauratostoma*, it is longest (35 – 38%). Here length of any body parts belong to two different genera may be same if it is presented in respect to total length, but must differ in respect to body length excluding tail. So it is better to calculate body parts in percentage of standard length i.e. body length excluding tail length (as used in morphometric description of a fish) and thus genus or species can be identified easily. Egg size is also species specific and often depends on body width rather than length. So it is presented in percentage of maximum body width.

Rao (1958) placed the genera *Travassosinema, Indiana* and *Pulchrocephala* under the family, Travassosinematidae on the basis of striking similarity of cephalic structure. Later Adamson and Van Waerebeke (1992) also supported the proposition though last two genera are parasites of mole cricket and *Travassosinema* is a parasite of millipede. However, a few authors namely Kloss, 1960 and Poinar, 1973 placed *Travassosinema* under family Thelastomatidae Travassos, 1920 and the other two umbraculum bearing genera, *Indiana* and *Pulchrocephala* in the family Pulchrocephalidae Kloss, 1959. Though cephalic umbraculum in all three genera are homologous structure (Adamson and Van Waerebeke, 1992), *Travassosiinema* differs from other two genera on the basis of three important characters: i). male of *Travassosinema* bears spicule which is absent in other two genera (Rao, 1958). ii). Eggs of *Indiana* and *Pulchrocephala* bear polar filaments which are absent in *Travassosinema* (Rao, 1958). iii). the origin of alae are different in these three genera. In *Travassosinema*, these are derived from fused adjacent lips in a six-lipped form but in other two genera, these are derived from three labia and three interlabia. Moreover, in Indiana, there are six additional alae (Adamson, 1987). Furthermore, the genera *Indiana* and *Pulchrocephala* are more similar to the other genera, parasite to mole cricket namely *Binema, Chitwoodiella, Isobinema* etc. On the basis of above mentioned morphometric dissimilarities, habitat and molecular data as studied by Morffe and Hasegawa (2017), only *Travassosinema* should be placed in the family Travassosinematidae. It requires further molecular study on the genera, *Indiana* and *Pulchrocephala* for confirmation.

## Acknowledgement

I am deeply indebted to my son Dr. Soumedranath Bhakat, Lund University, Sweden for his inspiration to concentrate my work on millipede and my wife Mrs. Mallika Bhakat for her mental support and boost in this pandemic phase. Finally I am grateful to all of my colleagues of Rampurhat College.

